# Odorization of Natural Gas: What are the Challenges?

**DOI:** 10.1101/2021.07.10.450231

**Authors:** Paul Wise, Steven Rowe, Pamela Dalton

## Abstract

Modern natural gas (NG) has little or no odor, so other compounds, usually mercaptans and thiols, are added as warning odorants. Federal regulations state that NG must be odorized so that it is readily detectable by people with normal senses of smell at one fifth the lower explosive limit, but regulations don’t define “readily detectable” or “normal senses of smell.” Methods to measure human odor detection have been available for decades. However, most previous work on NG odorants has underestimated human sensitivity, and measurements need to be repeated using the latest methods. More work is also needed to determine how odor sensitivity measured under optimal laboratory conditions is affected by real-world factors such as distraction and exposure to other odors in the environment. Regarding a “normal sense of smell,” healthy people vary over orders of magnitude in the concentrations they can detect, so samples of subjects should be chosen to reflect the range of differences in the population.

## Introduction

In the past, coal gas had a strong odor, primarily due to sulfur compounds (Robertson, 1980). Modern natural gas (NG), almost entirely formed of methane and ethane, has little or no odor, so sulfides and mercaptans are added to warn people of potentially dangerous leaks (Tenkrat, 2011). According to Federal regulations (49 CFR 192.625), “A combustible gas in a distribution line must contain a natural odorant or be odorized so that at a concentration in air of one-fifth of the lower explosive limit (LEL) [1% by volume in air for NG], the gas is readily detectable by a person with a normal sense of smell.” Many state regulations are similar. However, regulations do not specify the exact meaning of “readily detectable” or “a normal senses of smell,” nor do regulations specify exactly the types or amounts of odorant to add. Individual natural gas providers have devised working solutions based on their gas sources and infrastructure, with periodic checks on odor levels performed under field conditions by technical staff (Tenkrat, 2010). These solutions have proven effective in many cases since people often detect NG leaks by smell (Rawson et al., 2011).

How can odor research help? Toward clearer guidelines on what is “readily detectable,” researchers have suggested the general approach of measuring detectability of NG odorants under optimum (controlled laboratory) conditions, then applying a safety factor to account for real-world conditions.

### How do we measure odor detectability in the laboratory?

The two most common measures of odor detectability are detection thresholds and recognition thresholds. Detection threshold is the minimum odorant concentration observers can reliably discriminate from clean air. Detection, in a laboratory test scenario, is measured by presenting puffs of air to a panel of smellers. A smelling session consists of a number of “trials” in which a smeller samples one or more odorized puffs interspersed with one or more odorless puffs. The smeller must determine which puff(s) smell stronger, guessing if uncertain. The probability of correctly identifying the odorized puff(s) in a given trial increases as an s-shaped function of odorant concentration. One common definition of threshold is the concentration corresponding to detection performance half-way between chance (no better than guessing) and 100% correct. Abbreviated methods exist to estimate threshold concentration without having to measure the whole function. As in most endeavors, short-cuts entail compromise. Full detection functions (viz., percent correct at every concentration presented) provide the most reliable and precise data. Regardless, at detection threshold, a person can often determine that an odor is present, but may not be able to determine the source or significance of the odor.

Recognition thresholds, typically 2- to 10-fold higher than detection thresholds, are the minimum concentration needed to recognize the quality or identity of an odor (Hellman and Small, 1974). In some experimental methods, testers first present a fairly strong puff of odor as an example, then present lower concentrations to determine the lowest concentration which smells like the example. In other methods, smellers apply labels to a series of puffs to find the minimum concentration which receives an expected label (e.g., “gassy”). We can directly observe whether a smeller is correct or incorrect regarding which of a series of puffs is odorized (detection), but cannot directly observe whether an odor puff smells “gassy.” Thus, many recognition methods include “catch trials,” e.g., puffs of clean air or another odor to determine if the smeller is biased toward falsely reporting a puff smells “gassy.”

The concepts outlined above have long been understood in the study of human sensation and perception more generally, suggesting that determining a concentration that is “readily detectable,” at least in the laboratory, would seem easy to solve. However, even in controlled laboratory studies, measured thresholds for a given compound can vary, often up to a thousand-fold underscoring the fact that methodological details matter considerably (Devos et al., 1990).

Over the last several decades, researchers have made a concerted effort to understand which methodological details are crucial for measuring reliable and accurate odor thresholds, though the majority of these efforts were focused more on measuring the detectabilitiy of environmental odors more generally rather than gas odorization specifically (van Harreveld, Heeres, and Harssema, 1999; Mannebeck and Mannebeck, 2001; McGinley and McGinley, 2001). This work has resulted in new European Union standards (EN13725; CEN, 2003) that are gaining traction worldwide, though some aspects of EN13725, including how panelists are selected for testing, might not be optimal for the needs of the NG industry (a point to which we will return). As one might expect, good measurement depends in large part on a laboratory with clean air, a device (called an olfactometer) that allows precise and repeatable control of odor concentration, and puffs of sufficient flow rate and duration to support natural sniffing. For example, published thresholds for acetic acid, a key component of vinegar, span a ~1,000-fold range (Devos et al., 1990), but thresholds measured in three studies using precise olfactometry spanned only a 1.6-fold range and clustered at the lower end of the range of previously reported values (Cometto-Muñiz and Abraham, 2010). In brief, many of the historical reported odor thresholds have greatly underestimated the sensitivity of the population.

We expect to observe similar results with natural gas odorizers, and have begun an effort to update the extant data-base on sensitivity to NG odorants using rigorous methods and well-validated protocols. Initial results for two commonly used NG odorants, t-butyl mercaptan (TBM) and tetrahydrothiophene (THT), appear in Figure 1 (see appendix for methods). As predicted, our measured thresholds fell toward the lower end of the range of previously reported values (Figure 2). Thus, averages of reported values for NG odorants probably under-estimate sensitivity. The ongoing effort to update thresholds for NG odorants with state-of-the-art methods will provide an important base on which to build.

**Figure 1.**
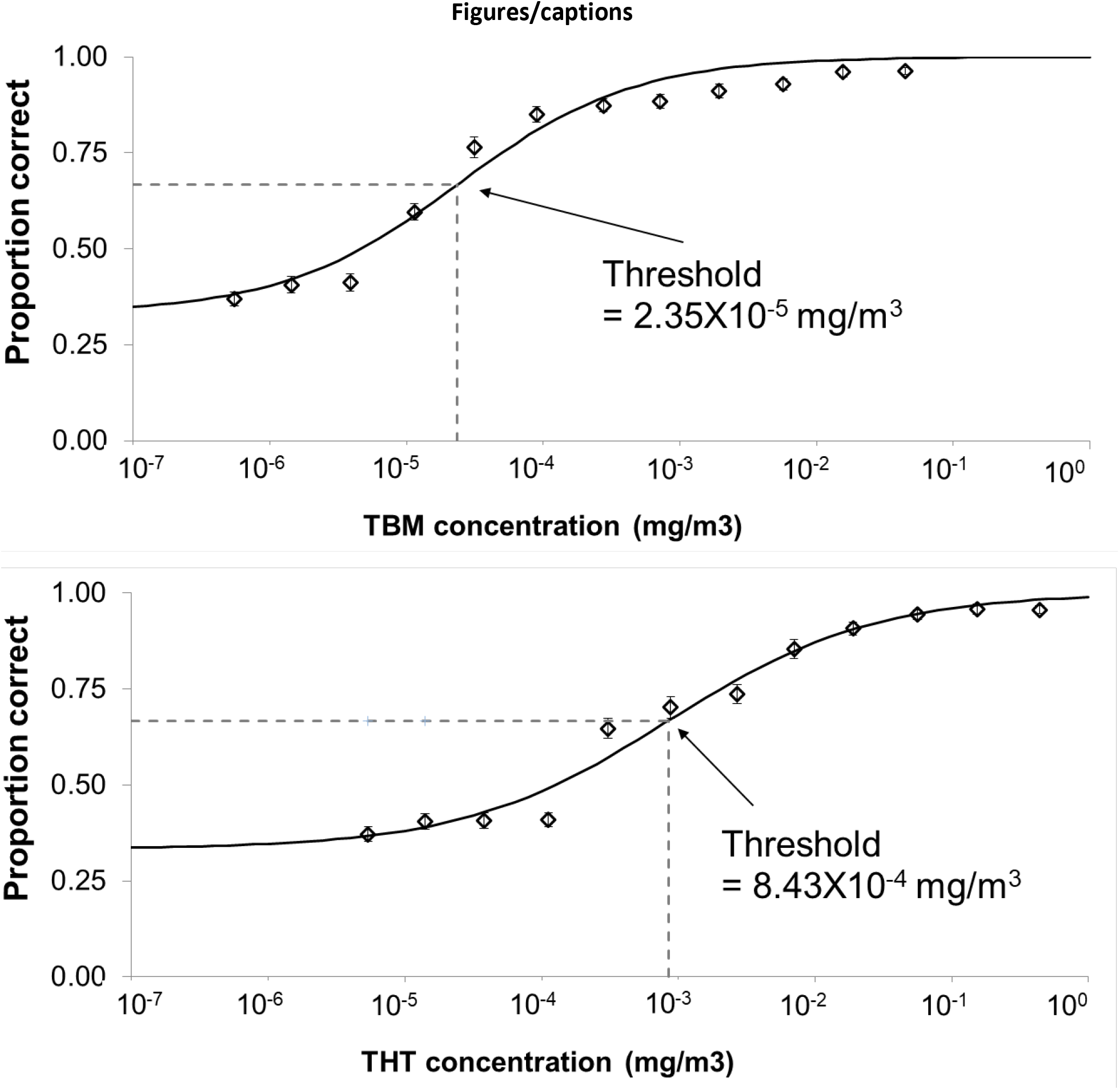
Odor detection functions for the common natural gas odorants t-butyl mercaptan (TBM) and tetrahydrothiophene (THT) measured using rigorous, air-dilution olfactometery. A panel of 41 subjects sniffed many sets of three odor puffs. One puff in each set of three puffs was odorized (target). The other two puffs were clean air (blanks). Subjects attempted to determine which puff smelled strongest, guessing if uncertain. The target puff spanned a range of concentrations from those subjects had no ability to detect (33.3% correct, guessing level) to those subjects could reliably detect. Detection threshold is typically defined as the odorant concentration corresponding to percent correct half way between guessing and perfect detection (in this case, 66.7%). Note that the large difference between TBM and THT suggests that detection of equal mass mixtures (like SCENTINEL^®^ T-50 from Chevron-Phillips) is driven largely by detection of TBM. Work on binary mixtures (data not shown) were consistent with this idea.

**Figure 2.**
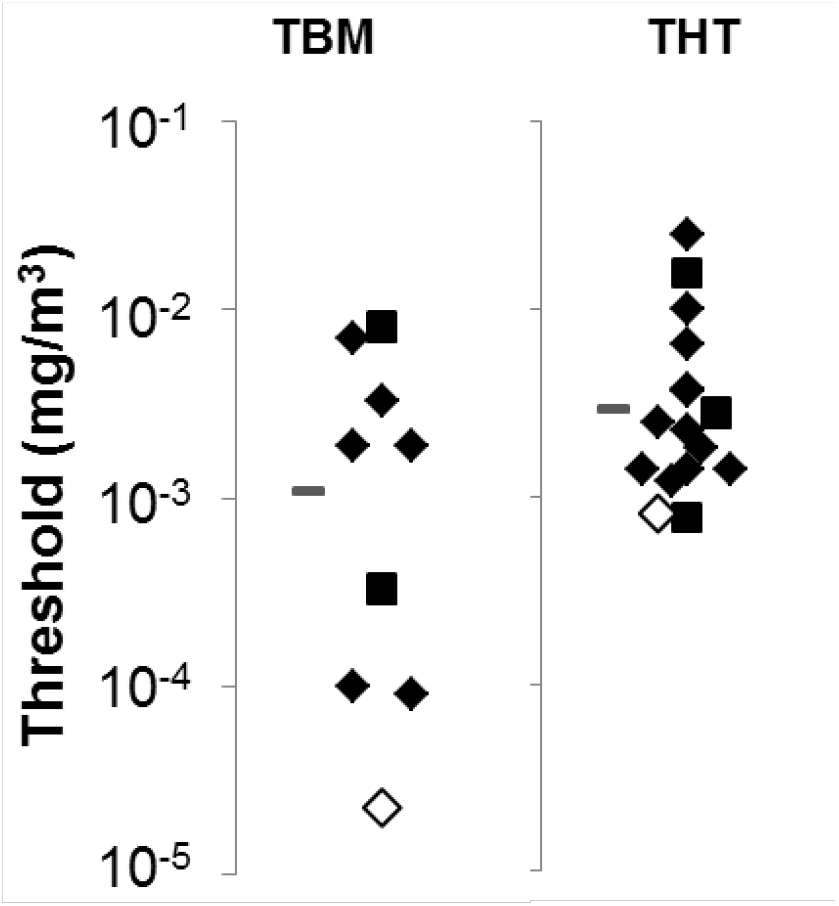
Distribution of previously measured odor thresholds for two commonly used natural gas odorizing agents, t-butyl mercaptan (TBM) and tetrahydrothiophene (THT). Filled diamonds represent previously measured detection thresholds. Filled squares represent previously measured recognition thresholds (see text for the difference between detection and recognition). Note that reported values cover a wide range of concentrations. Open diamonds represent detection thresholds we recently measured using state-of-the art methods (see Figure 1). Our measurements suggest that the average of previously reported values (grey hash marks) underestimate human sensitivity to these odorants.

### Factors which affect odor detection in the real-world

In the lab, panelists are engaged to actively detect odors with few distractions. People at home and work focus on their activities. Few studies have addressed the effect of attention and other cognitive factors on detection of NG odorants. One study found that people who were asked to read a message under low light were less likely to notice NG odorants in a laboratory than people who were not so distracted (Whisman et al., 1978). In another study, participants entered a travel trailer odorized with NG odorants, ostensibly to perform a consumer evaluation (Moschandreas et al., 1984). Despite various distractions, the proportion of participants who noticed gas odor was comparable to the proportion of participants who reached recognition threshold at a given concentration in controlled laboratory experiments. In short, evidence on distraction is mixed.

Adaptation is another potentially important factor. After steady or periodic exposure to a given odor, that odor tends to becomes more difficult to detect. Odor adaptation is often distinguished from habituation, in which a person stops noticing a smell but can detect it if they focus (related to the role of attention). In contrast, olfactory adaptation renders an individual incapable of smelling an odor without a period of recovery away from the odor. Regardless, adaptation to sulfides and mercaptans has been documented (Berglund et al., 1978; Moschandreas et al., 1984; Matsubasa et al., 2016). Adaptation can be particularly problematic when combined with the gradual fashion in which gas often builds up from leaks in indoor environments (Klusman, 1980; Larocque, 1981). Researchers have described comparable adaptation in laboratory studies, such that participants never detect a warning odor, even after it grows to relatively high levels (Wilby, 1990). However, the phenomenon is not well characterized in published literature.

Another potential issue is that, whereas olfactory laboratories have clean air by design, in daily life NG warning odors will be one of hundreds or thousands of odorous compounds in the environment. In general, odorous compounds tend to be mutually suppressive, such that any particular aroma will be less perceptible in a mixture (Thomas-Danguin 2014). There are numerous reports of NG which carries usual amounts of odorizing agents according to analytical measurements, but seems to have either little or no odor, or does not smell “gassy” (Bruno, 2009). This phenomenon remains unexplained, but other odors in gas, pipelines, or ambient air could suppress the perception of odorizing agents. A more common concern is that sulfur compounds, including some used in NG, are frequently encountered in daily life (e.g., odors of decay, odors from cooking meats and vegetables). This odor background could make NG odors more difficult to detect and recognize, with related problems caused by adaptation/habituation through constant exposure, though more work would be required to estimate how important and how large such effects might be.

Detection of NG warning odors in daily life require higher concentrations than the odor threshold concentrations determined in the lab. Our future studies should be conducted under conditions as realistic as possible, ideally: 1) in model dwellings and workspaces, 2) with distracted participants unaware that NG warning odor might be introduced, 3) with and without realistic odor backgrounds, and 4) with odorants entering and diffusing through the space in a realistic fashion and building up at varying rates (Hill and Pool, 1998).

### What is a “normal sense of smell?”

According to a large data-base, the most sensitive 5% among people aged 16 to 35 have odor thresholds about 100-fold lower than the least sensitive 5% (Hummel et al., 2007). For people aged 36-55, thresholds for the least and most sensitive differ by about 1000-fold. Variation has been on the same order of magnitude in smaller studies of “normal, healthy” people, including previous work on NG odorants (e.g., Wilby, 1969; Stevens et al., 1988; Figure 3). In addition, many people with a generally sensitive nose are insensitive to one or more particular odors (Amoore, 1968). At least some cases of “specific anosmia” (specific inability to smell) can be explained by genetically determined differences in smell receptors in the nose, analogous to color blindness in some respects (Knaapila et al., 2012; Logan, 2014). To the best of our knowledge, there have been no definitive demonstrations of specific anosmias to NG odorants. However, one study suggests that some people with good sensitivity to many sulfides and mercaptans prove relatively insensitive to particular NG compounds (Wilby, 1969).

**Figure 3.**
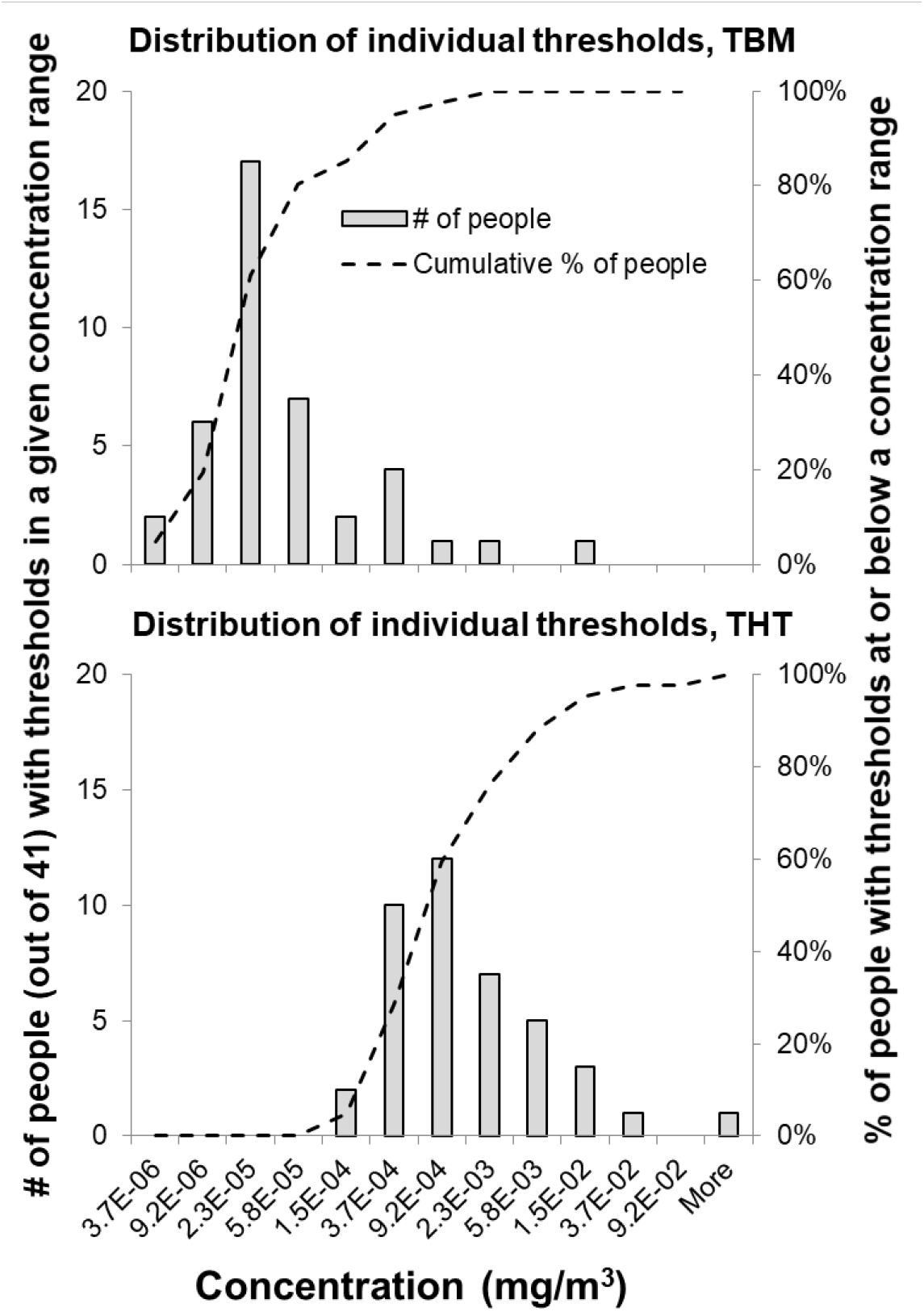
Thresholds calculated from data aggregated across individuals (as depicted in Figure 2) fail to convey important information, viz. that values for individuals spanned a range of about 1450-fold for t-butyl mercaptan (TBM), or about 230-fold excluding one extremely insensitive smeller. For tetrahydrothiophene (THT), values spanned a range of about 4900-fold (or about 770-fold excluding the same insensitive smeller. Smellers were deliberately selected to span a wide range ages (XX to XX) and olfactory abilities to be representative of healthy adults. This practice differs from EN13725, which specifies small samples chosen to be highly homogenous in sensitivity. Thus, EN13725 compliant methods without this modification do not provide estimates of the percentage of healthy adults able to detect an odor at a given concentration (dashed curves).

In addition to “normal” variation, up to 20% of people have olfactory dysfunction, with 4-6% being unable to smell at all (Brameson et al., 2004; Huttenbrink et al, 2013; Karpa et al., 2014; Mullol et al., 2012). Impaired individuals are often unaware of their deficit (Nordin et al., 1994; Murphy et al., 2002; Santos et al., 2004; Shu et al., 2011). Several common factors and conditions are associated with olfactory loss (Doty, 2009; Doty, 2015; Godoy et al., 2015; Ajmani et al., 2016; Rombaux et al., 2016), but the most common is aging (Attems et al, 2014; Doty and Karmath, 2014). In the United States, up to about 50% of people between the ages 65 and 80 and 75% of people over 80 have some dysfunction (Doty et al., 1984; Duffy et al., 1995; Murphy et al., 2002; Schubert et al., 2012). To make matters worse, the elderly tend to adapt to odors more readily and recover more slowly (Stevens et al., 1989), and often prove particularly insensitive to sulfur compounds used as NG odorizing agents (Chalke et al., 1958; Stevens et al., 1987; Wysocki and Gilbert, 1989; Cain et al., 1993; Larsson et al., 2000).

To calculate concentrations that offer the greatest benefit to the most people requires accurate data on the range of individual sensitivity across the population. Such data might also be useful for informing applicable regulations. Thus, our approach to updating the data-base of thresholds for NG odorants includes panelists with a wide range of sensitivities (Figure 3), tested by measuring full detection (Figure 1). Full functions provide much more reliable data on individuals than do the short-cut methods used in most past studies. Note that panels of individuals who are more representative of the general population is a departure from the guidelines in EN13725, which specifies testing small samples (6-8 panelists) who fall within a very narrow range of olfactory abilities. The EN13725 method of selecting panelists provides efficiency, but no information on the range of sensitivity across the general population for whom NG odors must serve as warning signals.

### Conclusions

Regulations require providers to odorize NG so that it is readily detectable by people with a normal sense of smell at one fifth the LEL, but regulations offer no guidelines regarding the meaning of “readily detectable” or “normal sense of smell.” Researchers have provided partial answers by measuring detection of NG odorizing agents under (optimal) laboratory conditions and suggesting that a safety factor will need to be applied for real-world conditions. Based on the data we report here, we find that extant data-bases of NG odor thresholds may under-estimate the sensitivity of the population. Thus, updating these values using modern methods, as we have begun, can provide a robust database for the industry’s needs.

## Appendix (methodological details for the threshold measurements reported in Figure 1)

Forty-one generally healthy adults (21 men, 20 women) aged 19 to 77 (mean=36.6) participated. Panelists were screened for smell sensitivity using the Sniffin’ Sticks™ butanol threshold test, a standardized test supported by a large, normative data-base (Oleszkiewicz et al., 10.1007/s00405-018-5248-1). The sample purposefully included people who spanned the full “normal” range, including the bottom 5^th^ percentile, the top 5^th^ percentile, and people of intermediate sensitivity. Procedures were conducted according to the principles in the Declaration of Helsinki and approved by an Institutional Review Board (IRB) at the University of Pennsylvania. Subjects provided written, informed consent using IRB approved forms before any experimental procedures were conducted.

Odor stimuli included tert-butyl mercaptan (TBM; CAS# 75-66-1) and tetrahydrothiophene (THT; CAS# 110-01-0). Both compounds have been widely used as natural gas warning odors, and odor thresholds for both have been reported by multiple groups (Devos et al., 1990; Tenkrat, 2011). The compounds were purchased in compressed cylinders (custom made; Scott Specialty Gas, Philadelphia PA). According to the manufacturer, the TBM cylinder contained 0.0762 g of TBM in 4732.98 g nitrogen (5 ppm by volume, or 20.14 mg/m^3^). The THT cylinder contained 0.746 g of THT in 4742.5 g nitrogen (50 ppm by volume, or 196.78 mg/m^3^). These concentrations were verified using gas chromatography/mass spectrometry. The cylinders were used to fill Tedlar™ gas sampling bags, which in turn served as odor-sources for an air-dilution olfactometer (TO Evolution, Olfasense™, Kiel, Germany) compliant with EU standards for precision olfactometry (EN13725; CEN, 2003). The other olfactometer input (used to dilute the contents of the sample bags and for blanks) was triple-filtered, medical-grade air. Twelve dilutions of each compound were presented (~2.8-fold steps), ranging from 5.50 × 10^−7^ to 4.49 × 10^−2^ mg/m^3^ for TBM and 5.38 × 10^−6^ to 4.38 × 10^−1^ mg/m^3^ for THT. Olfactometer dilutions were factory calibrated and certified, and proportionality was verified at the output of the olfactometer using gas chromatography.

Threshold tests were conducted in a clean environmental chamber with computer controlled temperature and humidity, with a ventilation rate of more than 17 m^3^/min. Subjects sampled stimuli from a sniffing cone, with a flow rate of 20 l/min. The task was temporal, 3-alterantive forced-choice (3-AFC), with one odorized air pulse randomly interspersed with two clean air pulses (6 s per pulse). Subjects indicated which of the three pulses smelled strongest, guessing if uncertain, by pressing virtual buttons on a touch pad. Subjects completed blocks of 12 trials, or each concentration of a particular compound in ascending order of concentration, with 40 s between trials. Subjects completed two ascending blocks for each odorant in a session, with pauses of ~5 min between blocks. After a practice session, subjects completed 9 more sessions, for up to 18 trials per concentration and stimulus (one subject completed 13 trials per concentration, and there were occasional missing data for others due to failure to enter responses).

For each stimulus (and for each individual subject), percent correct was plotted vs. concentration, and detection curves were fit with cumulative logistic (Hill) functions. Lower asymptotes were fixed at 33% (chance for a 3-AFC task). Upper asymptotes were fixed at 100%. Slope and point of inflection (i.e., threshold) were free parameters. Functions were fit *via* least-squares regression using the robust Solver^®^ function in Microsoft Excel (Version 14.0.7212.5000)(Gadagkar & Call, 2015). Fits for individual subjects were reasonable for the number of trials collected: average r^2^ (proportion of variance for which fitted functions account) was 0.88 (standard deviation of 0.10).

